# Drosophila *small ovary* gene ensures germline stem cell maintenance and differentiation by silencing transposons and organising heterochromatin

**DOI:** 10.1101/383265

**Authors:** Ferenc Jankovics, Melinda Bence, Rita Sinka, Anikó Faragó, László Bodai, Aladár Pettkó-Szandtner, Karam Ibrahim, Zsanett Takács, Alexandra Brigitta Szarka-Kovács, Miklós Erdélyi

**Affiliations:** Institute of Genetics, Biological Research Centre of the Hungarian Academy of Sciences, Szeged, Hungary; Department of Genetics, University of Szeged, Szeged, Hungary; Department of Biochemistry and Molecular Biology, University of Szeged, Szeged, Hungary; Institute of Plant Biology, Biological Research Centre of the Hungarian Academy of Sciences, Szeged, Hungary

**Keywords:** Drosophila, stem cell niche, piRNA, chromatin, HP1a

## Abstract

Self-renewal and differentiation of stem cells is one of the fundamental biological phenomena relying on proper chromatin organisation. In our study, we describe a novel chromatin regulator encoded by the *Drosophila small ovary (sov)* gene. We demonstrate that *sov* is required in both the germline stem cells (GSCs) and the surrounding somatic niche cells to ensure GSC survival and differentiation. *Sov* maintains niche integrity and function by repressing transposon mobility, not only in the germline, but also in the soma. Protein interactome analysis of Sov revealed a physical interaction between Sov and HP1a. In the germ cell nuclei, Sov co-localises with HP1a, suggesting that Sov affects transposon repression as a component of the heterochromatin. In a position effect variegation assay, we found a dominant genetic interaction between *sov* and HP1a, indicating their functional cooperation in promoting the spread of heterochromatin. An *in vivo* tethering assay and FRAP analysis revealed that Sov enhances heterochromatin formation by supporting the recruitment of HP1a to the chromatin. We propose a model in which *sov* maintains GSC niche integrity by regulating piRNA-mediated transposon silencing as a heterochromatin regulator.

**Summary statement:** *Small ovary* maintains the integrity of the stem cell niche by regulating piRNA-mediated transposon silencing acting as a key component of the heterochromatin.

## Introduction

The eukaryotic genome is organized into structurally distinct and functionally specialized chromatin domains, called euchromatin and heterochromatin. The euchromatic domain contains actively transcribed genes, whereas the heterochromatin is mainly associated with a repressive transcriptional state. The heterochromatin is enriched in repetitive elements and transposons which occupy the centromeric and telomeric regions of the chromosomes. Another domain of the heterochromatin is formed at regulatory regions of genes that have to be transcriptionally repressed at specific stages of development. The heterochromatin is epigenetically defined by a combination of specific covalent modifications of histone molecules. The formation of the heterochromatin is accompanied by trimethylation of Histon3 at Lysin9 (H3K9me3) which recruits Heterochromatin protein 1a (HP1a, encoded by Su(var)205) initiating the formation of the repressive chromatin environment. Heterochromatic domains are organized around HP1 into phase-separated liquid compartments that physically compact chromatin and recruit additional repressive components (Larson et al., 2017; Strom et al., 2017). Kinetic analysis of HP1-chromatin binding revealed a complex interaction between heterochromatin components; however, the precise molecular mechanisms required for formation and maintenance of heterochromatin domains are not completely understood (Bryan et al., 2017).

In eukaryotes, a heterochromatin-dependent, small non-coding RNA-based defense system has been evolved against transposon-induced mutagenesis (Tóth et al., 2016). Central components of this pathway are the Piwi-interacting RNAs (piRNAs). Long precursors of the piRNAs are transcribed from both uni-strand and dual-strand piRNA clusters containing transposon sequences. Following piRNA biogenesis in the cytoplasm, mature short piRNAs associate with members of the Piwi class of the Argonaute protein superfamily (Piwi, Aub and Ago3 in Drosophila) forming RISC complexes. In the germ cells, Aub and Ago3-RISC complexes mediate the post-transcriptional silencing of the transposons by inducing the degradation of transposon transcripts in the cytoplasm. The Piwi-RISC complex, however, migrates into the nucleus and inhibits transposon transcription. In the somatic cells of the ovary, exclusively the Piwi-RISC mediated transcriptional silencing inhibits transposon activity.

Two steps of the piRNA pathway have been shown to depend on heterochromatin function. First, long precursors of the piRNAs are transcribed from piRNA clusters located mainly at the heterochromatic regions of the genome. Disruption of heterochromatin formation by *eggless/dSetdbl (egg)* or *HP1a* mutations impedes the transcription of the clusters resulting in derepression of transposons (Rangan et al., 2011; Teo et al., 2018). The second heterochromatin dependent step of the piRNA-pathway is the transcriptional silencing of the transposon transcription. At the transposon loci, Piwi-RISC inhibits transposon transcription by inducing the formation of a repressive heterochromatic environment on the transposon loci. Transcriptional silencing of the transposons includes the deposition of repressive H3K9me3 modification marks and the recruitment of HP1a to the chromatin of the transposon locus.

The Drosophila oogenesis provides an excellent model for understanding mechanisms of heterochromatin formation and its function in gene expression regulation and transposon silencing. In the ovary, repeated divisions of germline stem cells (GSCs) ensures continuous production of germ cells (Eliazer and Buszczak, 2011). GSCs reside stem cell niches which are located in the germaria at the anterior tip of the ovary. The GSC niches are composed of three somatic cell types, terminal filament cells, cap cells and escort cells (ECs), which provide physical and signaling milieu required for GSC self-renewal and differentiation. Mitotic division of the GSC reproduces the GSC and generates a committed progenitor cell, the so-called cystoblast. The cystoblast has a limited division capacity and generates 16 interconnected cyst cells. One of the cyst cells differentiates into an oocyte whereas the remaining 15 cyst cells become supportive nurse cells. The developing germ cells are surrounded by an epithelial monolayer of somatic follicle cells forming an egg chamber.

In a previous RNAi-based screen for genes regulating germ cell behavior, we have identified several essential chromatin regulators, such as *Su(var)205* and *Su(var)2-10,* to be involved in germ cell development (Jankovics et al., 2014). In the same screen, we identified the annotated CG14438 gene which has been shown to be involved in transposon silencing and to co-immunoprecipitate with HP1a (Alekseyenko et al., 2014; Czech et al., 2013; Muerdter et al., 2013). To gain a better understanding of chromatin regulation during germ cell development we analyzed the function of CG14438 in *Drosophila* oogenesis. Here, we show that CG14438 is identical to *small ovary* (*sov*) gene and it is a novel chromatin regulator that promotes heterochromatin formation by stabilizing the association of HP1a with the chromatin. Our results suggest that Sov suppresses transposon activity by regulating the transcription of the dual strand piRNA clusters and components of the piRNA pathway. In the stem cell niche, *sov* function is required both in the somatic and in the germ cells to ensure GSC maintenance and differentiation.

## Results

### CG11438 and *small ovary (sov)* are identical

CG14438 encodes for a single large protein of 3313 amino acids (Fig.1A). The N-terminal half of the protein is highly unstructured and contains a putative intrinsically disordered RGG/RG domain mediating degenerate specificity in RNA binding (Ozdilek et al., 2017; Thandapani et al., 2013). The C-terminal half of the protein contains 21 zinc-finger domains and a PxVxL pentamer motive, a canonical HP1 binding domain (Smothers and Henikoff, 2000).

**Fig.1.**
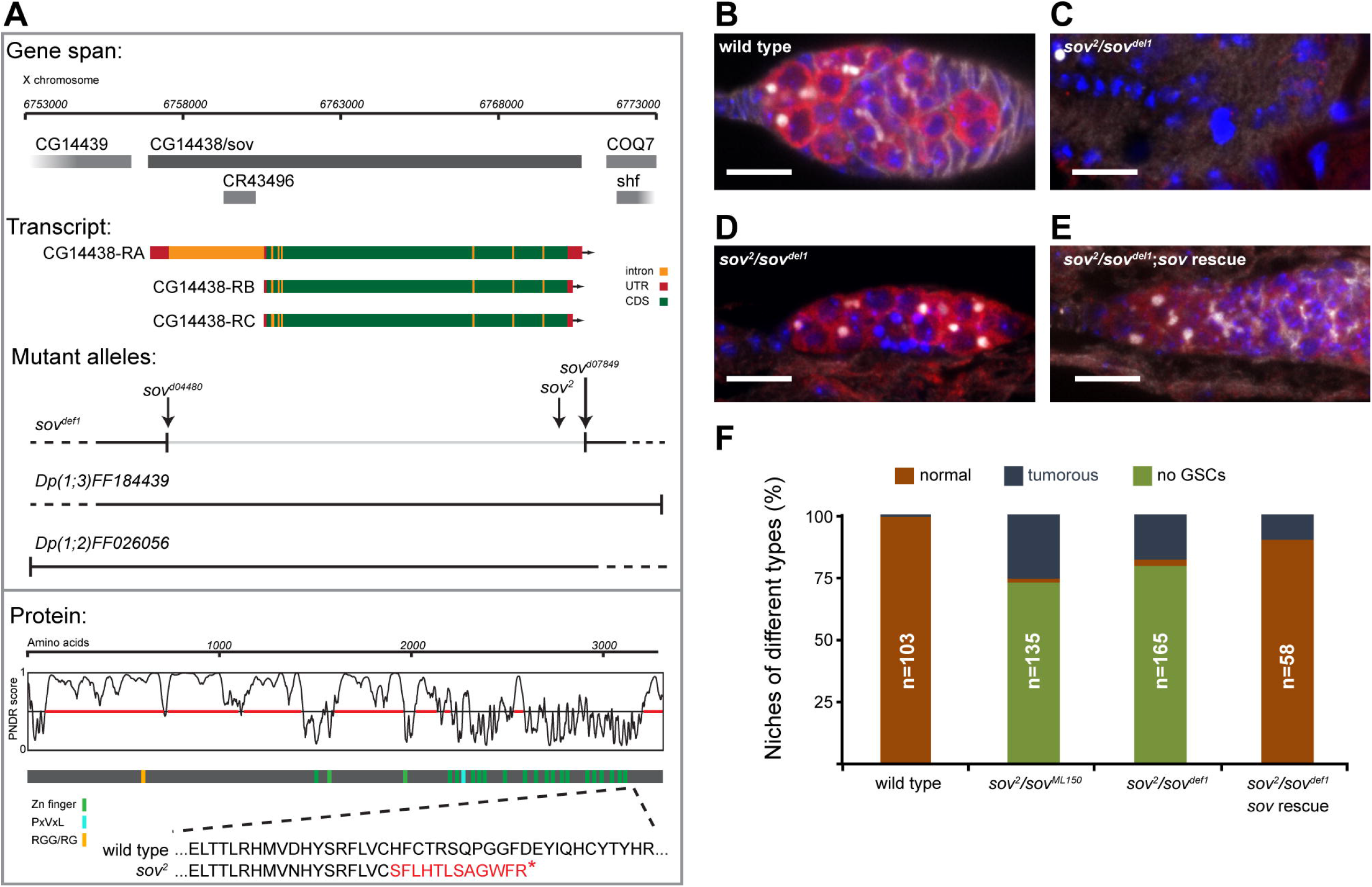
Sov mutations lead to GSC loss and formation of tumors. (Schematic representation of the *sov* (CG14438) locus showing *sov* transcripts and the mutant alleles used in this study. Domain composition and prediction of disordered regions of Sov protein is shown (Dunker et al., 2001). PNDR score >0.5 represents disorder and is indicated by red line. Alignment shows the wild type and *sov^2^* mutant Sov protein sequences. Red letters indicate additional amino acids generated by a frame-shift mutation in *sov^2^*. (B-E) Immunostaining of a wild type germarium (B), and a *sov^2^/sov^del1^* germarium with GSC loss (C) and with germ cell tumor (D). (E) Rescue of the niche defects in a *sov^2^/sov^del1^; Dp(1;2)FF194438* germarium. Spectrosomes and fusomes are labelled with HTS (white), germ cells are labelled with Vasa (red), DAPI is blue. Bars: 10 μm.(F) Quantification of the *sov* mutant phenotypes.

We generated a null allele of *CG14438* which removes the entire coding region of the gene (Fig.1A) (Ryder et al., 2004). Animals homozygous for the novel deletion allele, which we called *CG14438^del1^,* died at the third larval stage indicating that *CG14438* is an essential gene. Complementation analysis between *CG14438^dee1^* and alleles of genes mapping to the same genomic region revealed that *CG14438^de11^* does not complement alleles of *small ovaries (sov).* To confirm that *CG14438* corresponds to sov, we performed a series of rescue experiments. The Dp(1;3)DC486 duplication, which covers a 92.5 kb genomic region around the *CG14438/sov* locus rescued all *sov* phenotypes indicating that the *sov* gene is localized in this genomic region. Identical rescue was observed with two overlapping genomic transgenes (Dp(1;2)FF026056 and Dp(1;2)FF184439) (Fig.1A). Sequencing of the *sov^2^* allele revealed a point mutation in the *CG14438* coding region generating a premature stop codon after amino acid 3151 which results in a truncated mutant protein lacking 152 C-terminal amino acids. (Fig.1A) Taken together, the complementation analysis, the rescue experiments and the sequencing of the *sov* mutant allele indicate that CG14438 and *sov* are identical.

### Sov is required for GSC maintenance and differentiation

*Sov* is an essential gene and its hypomorphic allelic combinations result in similar ovarian morphology to that of *CG14438* RNAi (Jankovics et al., 2014; Wayne et al., 1995). To gain insights into the function of *sov* in germ cell development, we made use of the hypomorphic allelic combinations *sov^2^/sov^ML150^* and *sov^2^/sov^dee1^.* Females were sterile and exhibited rudimentary ovaries. In most of the germaria, no germ cells were found indicating a role for *sov* in germ cell maintenance (Fig.1C,F). In addition to the agametic germaria, ca. 30% of the mutant germaria exhibited germ cell tumors. (Fig.1D,F). Each tumor cell contained a single spectrosome, a hallmark of the GSC or cystoblast-like undifferentiated germ cells, indicating that *sov* is required not only for GSC maintenance but also for GSC differentiation. The GSC maintenance and differentiation defects were rescued by the Dp(1;2)FF184439 transgene expressing the *sov* gene from its genomic context (Fig.1E,F).

### *Sov* is required cell-autonomously for GSC maintenance at the adult stage

To narrow the temporal and spatial requirement of *sov* in germ cell development, tissuey-specific RNAi and clonal analysis were performed. Silencing of *sov* with the germline-specific nosGal4 driver caused female sterility. Young females possessed normal-looking ovaries (Fig.2B,D) and laid eggs which did not hatch. However, depletion of *sov* in the germ line resulted in a progressive loss of GSCs. In four-week old females, most of the niches lost the GSCs and contained no germ cells indicating that *sov* is required cell-autonomously in the germ line for GSC maintenance (Fig.2C,D).

**Fig.2.**
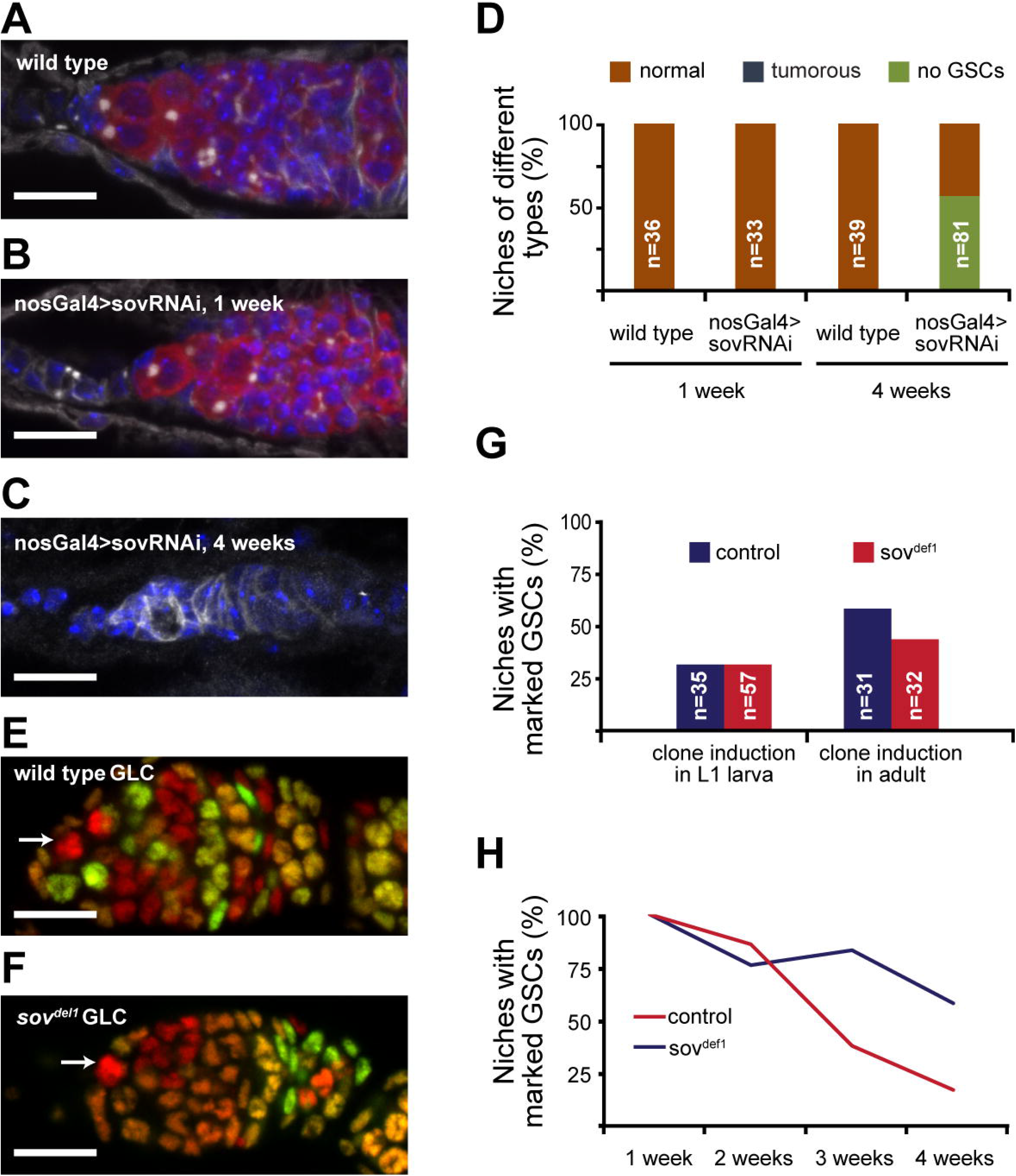
Sov is required cell-autonomously for GSC maintenance. (A-D) Germline-specific *sov* silencing. Immunostaining of a wild type germarium (A) and a germarium of a one-week old (B) and of a four-week old (C) *nosGal4>sovRNAi* female. Spectrosomes and fusomes are labelled with HTS (white), germ cells are labelled with Vasa (red), DAPI is blue. Bars: 10 μm. (D) Quantification of the phenotypes induced by germline specific *sov* silencing. (E-H) Analysis of germline clones (GLCs). (E-F) Immunostaining of a germarium with a wild type (E) and *sov^del1^* mutant (F) GLCs. Cells ubiquitously express Histon2Av:RFP (red) and Histon2Av:GFP (green). Having lost Histon2Av:GFP, GLCs appear red in the immunostaining (arrows). Bars: 10 μm. (G) The frequency of the induction of wild type and *sov^del1^* homozygous mutant GLCs. (H) Normalized frequency of germaria carrying wild type and *sov^del1^* GLCs at various time points after clone induction.

Cell-autonomous requirement of *sov* in germ cell maintenance was confirmed by analysis of *sov* mutant germline clones. Homozygous mutant *sov^del1^* germ cells were induced by FLP/FRT mediated recombination and identified by the lack of the GFP marker gene (Fig.2E,F). To analyze *sov* function at the larval stage, *sov* mutant germ line clones were induced in L1 larvae and the phenotype was analyzed in adults. In three-day old females, GSCs were found in the *sov* mutants similar to the wild type clones indicating that *sov* mutant larval germ cells can populate the niche and can develop into normal GSCs (Fig.2G). This indicates that *sov* function is dispensable in the germ line between L1 and the adult stage. To analyze *sov* function specifically in adult GSCs, clones were induced in the germ cells of young females. Both in the control and in the mutant niches, GFP-minus GSCs appeared in the first week after clone induction (ACI) (Fig.2G). Wild type control clones were maintained even after four weeks ACI; however, the number of niches carrying *sov* mutant GSCs decreased (Fig.2H).

Taken together, our data show that *sov* is required for GSC maintenance intrinsically in the germline at the adult stage. Remarkably, loss of *sov* in the GSCs located in niches composed of wild type somatic cells did not induce tumor formation, indicating that the differentiation defect observed in *sov* mutants is not germ-cell-autonomous.

### Sov is required in ECs for GSC maintenance, germ cell differentiation, and EC survival

Formation of tumors composed of undifferentiated GSC-like cells and GSC loss could be a consequence of defects in the somatic cells of the niche. Depending on their position in the germarium, ECs have two distinct functions (Wang and Page-McCaw, 2018). Anterior ECs promote GSC self-renewal and maintain GSCs in stem cell state. Posterior ECs, however, promote GSC differentiation. Thus, loss of ECs in *sov* mutants results in a dual phenotype: GSC loss if the anterior ECs are lost, or GSC tumor, if the posterior ECs die. In *sov* mutant niches, the number of the ECs was reduced indicating that *sov* is required for EC survival (Fig.3B,E). Consistent with the EC loss, we observed accumulation of activated Caspase3 in the *sov* mutant niches (Fig.S1). A similar reduction in EC numbers was observed when *sov* was silenced specifically in the ECs by the c587Gal4 driver, indicating that the requirement of *sov* for EC survival is cell autonomous (Fig.3C,D,E).

**Fig.3.**
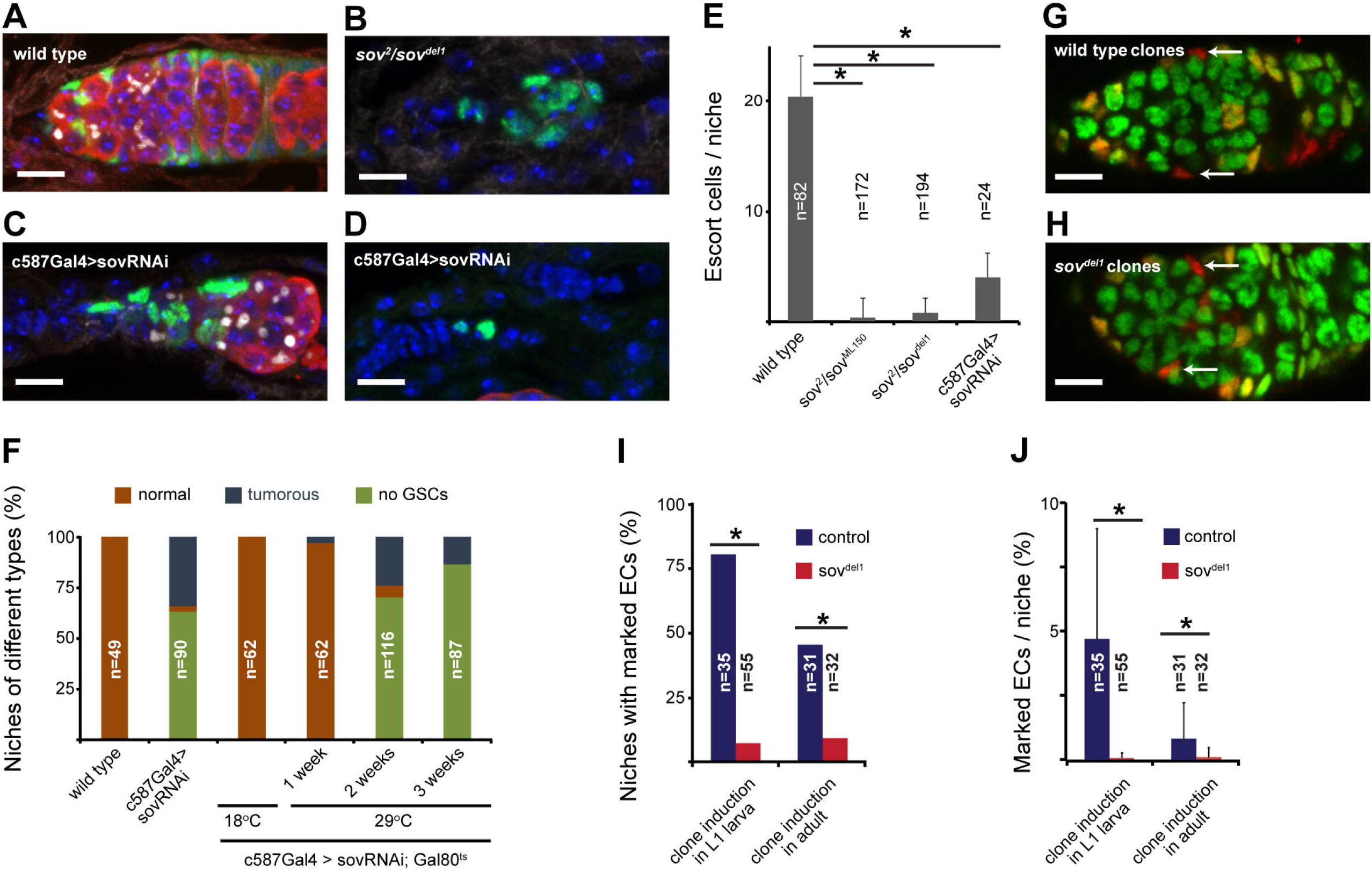
Sov is required in ECs for GSC maintenance, germ cell differentiation, and EC survival. (A-D) Immunostaining of a wild type germarium (A) and a *sov^2^/sov^del1^* germarium with reduced EC number (B). Immunostaining of *c587Gal4>sovRNAi* germaria with reduced EC number and exhibiting germ cell tumor (C) and GSC-loss (D) phenotypes. Spectrosomes and fusomes are labelled with HTS (white), germ cells are labelled with Vasa (red), ECs are labelled with Tj (green), DAPI is blue. Bars: 10 μm. (E) Quantification of EC number in *sov* mutant and *c587Gal4>sovRNAi* niches. Mean±s.d. is shown, T-test, * p<0.05. (F) Quantification of the mutant niche phenotypes. (G-J) Analysis of EC clones. (G,H) Immunostaining of a germarium with wild type (G) and *sov^del1^* mutant (H) EC clones. Cells ubiquitously express Histon2av:RFP (red) and Histon2av:GFP (green). EC clones that have lost Histon2av:GFP appear red in the immunostaining (arrows). Bars: 10 μm. (I,J) Mean±s.d. is shown, T-test, * p<0.05. (I) The frequency of the induction of wild type and *sov^del1^* homozygous EC clones. (J) Frequency of niches carrying wild type and *sov^del1^* EC clones.

Next, we investigated how somatic *sov* function affects germline behavior. Therefore, the c587Gal4 line was used to induce *sov* silencing in the ECs and the germ cells were analyzed. Silencing of *sov* in the ECs induced GSC loss and formation of GSC-like tumors in the niches (Fig.3C,D,F). This indicates that *sov* is required in the ECs in a non-cell autonomous manner for GSC maintenance and differentiation.

This *sov* RNAi phenotype in GSCs could be a consequence of earlier defects induced in the larval ancestors of the ECs. To induce adult-specific *sov* silencing in the ECs, the temperature-sensitive Gal80^ts^ mutant was used. At the permissive temperature (18°C), *sov* is not silenced in *c587Gal4;sovRNAi;Gal80^ts^* flies and no abnormalities are detectable in the niche (Fig.3F). However, shifting the adult females to the restrictive temperature (29°C) licensed Gal4 driven *sov* silencing in the ECs which, in turn, resulted in germline defects similar to those of *sov* mutants. Two weeks after RNAi induction, germ cell tumors were formed and GSCs were lost (Fig.3F).

Homozygous EC clones were induced in L1 larvae or in one-day old adult females, and the number of germaria containing homozygous *sov* mutant and wild type EC clones were determined in four-day old females (Fig.3G,H). The frequency of germaria containing *sov* mutant ECs was reduced compared to wild type clone frequency (Fig.3I). These germaria contained fewer *sov* mutant ECs, confirming the results obtained by analysis of mutant allelic combinations and the RNAi data on the requirement of *sov* in EC survival (Fig.3J).

In summary, *sov* is required for EC survival in the adult niches, which ensures GSC differentiation and maintenance in a non-cell-autonomous manner.

### Sov promotes GSC differentiation by restricting dpp-signalling activity in the niche

In the niche, a complex regulatory network controls GSC differentiation. To explore further the function of *sov* in GSC differentiation, we analyzed the activity of the signaling pathways involved in communication between different cell types of the niche.

In GSCs, the repression of *bam* expression prevents differentiation of the stem cells. In the wild type, *bam* expression is initiated in that GSC daughter cell that loses physical contact with the cap cells and adopts cystoblast fate. In *sov* mutant germaria, no *bam* expression was detected when monitored with the reporter line bam-GFP (Fig.S2A,B). Forced expression of *bam* from the heat shock inducible hs-bam transgene was sufficient to induce differentiation of the *sov* mutant germ cells, as indicated by the formation of fusome containing cysts and an almost complete lack of GSC-like tumors in the *sov^2^/sov^M^^L150^; hs-bam* females (Fig.S2C-E). Based on these data, we conclude that *sov* act upstream of *bam* in the GSC differentiation process.

In the niche, the primary factor that represses *bam* expression in the GSCs is Dpp, the Drosophila TGFβ homolog. In the wild type, Dpp activity is restricted to the GSCs and can be monitored by the nuclear translocation of pMad. Sov mutant tumor cells located outside of the GSC niche accumulate pMad in their nuclei, indicating that expanded Dpp activity is responsible for the maintenance of *bam* repression in the *sov* mutant germ cells (Fig.S2F,G). Restriction of Dpp activity to the GSCs can be adjusted by controlled diffusion of the secreted Dpp ligand. In wild type germaria, ECs extend long cellular protrusions that enwrap differentiating GSC daughter cells, separating them from the Dpp signal (Fig.S2H). Sov mutant ECs, however, fail to extend protrusions, indicating that *sov* controls the accessibility of the secreted Dpp (Fig.S2I). Niche abnormalities caused by the lack of EC protrusions resembles the phenotypes that have been observed in niches with impaired Wnt4 function. To test the involvement of *sov* in Wnt4 mediated niche regulation, Wnt4 expression was analyzed in *sov* mutants. RT-qPCR revealed that Wnt4 mRNA levels were reduced in *sov* mutant niches, indicating that *sov* is required for Wnt4 expression (Fig.S2J). Forced expression of Wnt4 in *sov* RNAi ECs by two different transgenic constructs, however, did not rescue the germ cell differentiation defects caused by *sov* silencing, indicating that *sov* regulates GSC development not exclusively by promoting Wnt4 expression (Fig.S2K).

Taken together, sov-mediated Wnt4 expression in the ECs promotes protrusion formation that separates the GSC daughter cell from the Dpp source and enables Bam-driven differentiation of the GSC daughter cell.

### Sov is required in both somatic and germline cells for transposon repression

Remarkably, GSC and EC defects of *sov* mutants resemble that of *egg, HP1a,* or *piwi,* essential regulators of heterochromatin formation and transposon repression in *Drosophila*. Thus, we hypothesized that Sov regulates niche integrity by suppressing transposon activity via heterochromatin regulation. To test this hypothesis, we first analyzed transposon expression in both the somatic and germ cells of the niche using transposon sensors, RT-qPCR and RNA-seq.

Transposon sensors contain transposon-derived sequences that are targeted by the piRNAs, resulting in repression of the LacZ reporter gene. First, we utilized the germline-specific *HET-A* and the *Burdock* transposon sensors (Dönertas et al., 2013). MTD-Gal4 driver was used to drive the germline-specific expression of three independent *sov* RNAi lines. In all *MTDGal4>sovRNAi* ovaries, a robust β-Galactosidase (β-Gal) expression was detected from both transposon sensors, suggesting derepression of transposons (Fig.4A-D). Consistent with these results a strong accumulation of the endogenous germline-dominant HET-A and Burdock transposon mRNA levels was detected in *nosGal4>sovRNAi* ovaries using RT-qPCR (Fig.4I). To determine the global effect of *sov* on the steady state RNA levels of all transposon classes, polyA-RNAs were sequenced from *nosGal4>sovRNAi* and *nosGal4>wRNAi* control ovaries (RNA-seq). In *sov* RNAi ovaries, a robust upregulation of transposon transcript levels was detected, indicating that *sov* is required for repression of transposon activity in the germline (Fig.4J).

**Fig. 4.**
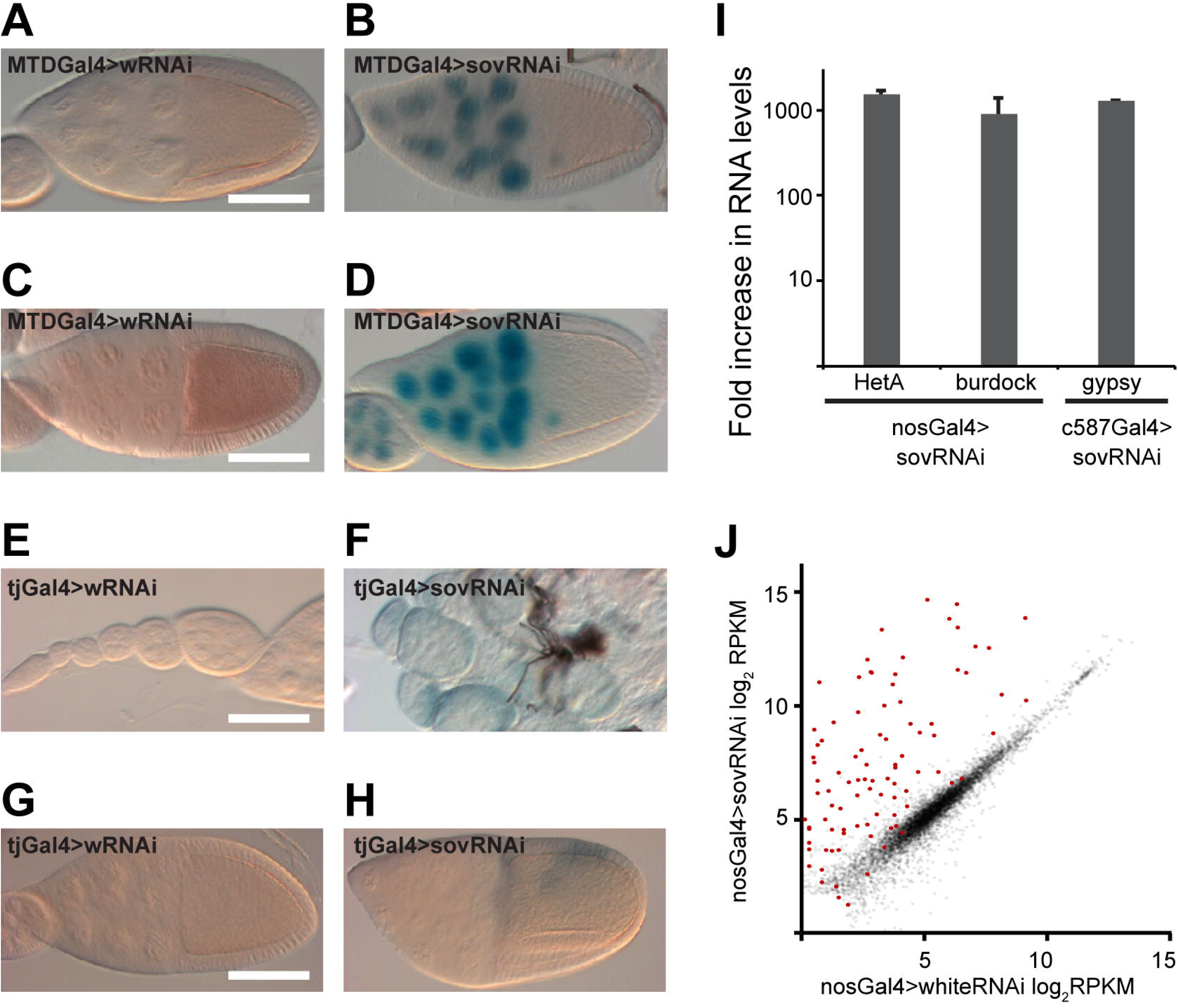
Sov is required in both somatic and germline cells for transposon repression. (A-H) Shown are β-Gal stainings of wild type and sov-silenced egg chambers expressing HeT-A (A,B) Burdock (C,D) gypsy (E-H) reporters. For *sov* silencing, TRiP-HMC04875 (H) and 3-2 (B,D,F) was used. (I) Displayed are fold changes in steady-state mRNA levels of indicated transposons measured by RT-qPCR. (J) Scatter plot showing expression levels of genes (gray) in *nosGal4>sovRNAi* ovaries. transposons are in shown in red.

To analyze the effect of *sov* on transposon silencing in the soma, we made use of the *gypsy-* LacZ transposon sensor (Dönertas et al., 2013). Silencing of *sov* with the 3-2 and KK103679 RNAi lines resulted in severe morphological abnormalities of the ovaries, which made the analysis of the *gypsy*-LacZ reporter difficult. Nevertheless, we detected a strong β-Gal accumulation by Tj-Gal4 driven expression of the silencing constructs, indicating transposon derepression by *sov* silencing (Fig.5E,F). Silencing of *sov* with the week HMC04875 RNAi line resulted in the formation of normal egg chambers with normal-looking somatic cells and the oogenesis was completed. Despite the modest phenotypic consequences of *sov* silencing by this RNAi construct, a weak β-Gal expression was detected in the somatic cells (Fig.4G,H). Results obtained through analysis of the gypsy-LacZ transposon sensor were confirmed by measuring endogenous *gypsy* mRNA levels in the *sov* mutant ovaries. RT-qPCR on c587Gal4*>sovRNAi* ovaries revealed a robust accumulation of the endogenous *gypsy* mRNAs, indicating that *sov* functions in the repression of the somatic transposon activity (Fig.4I).

**Fig. 5.**
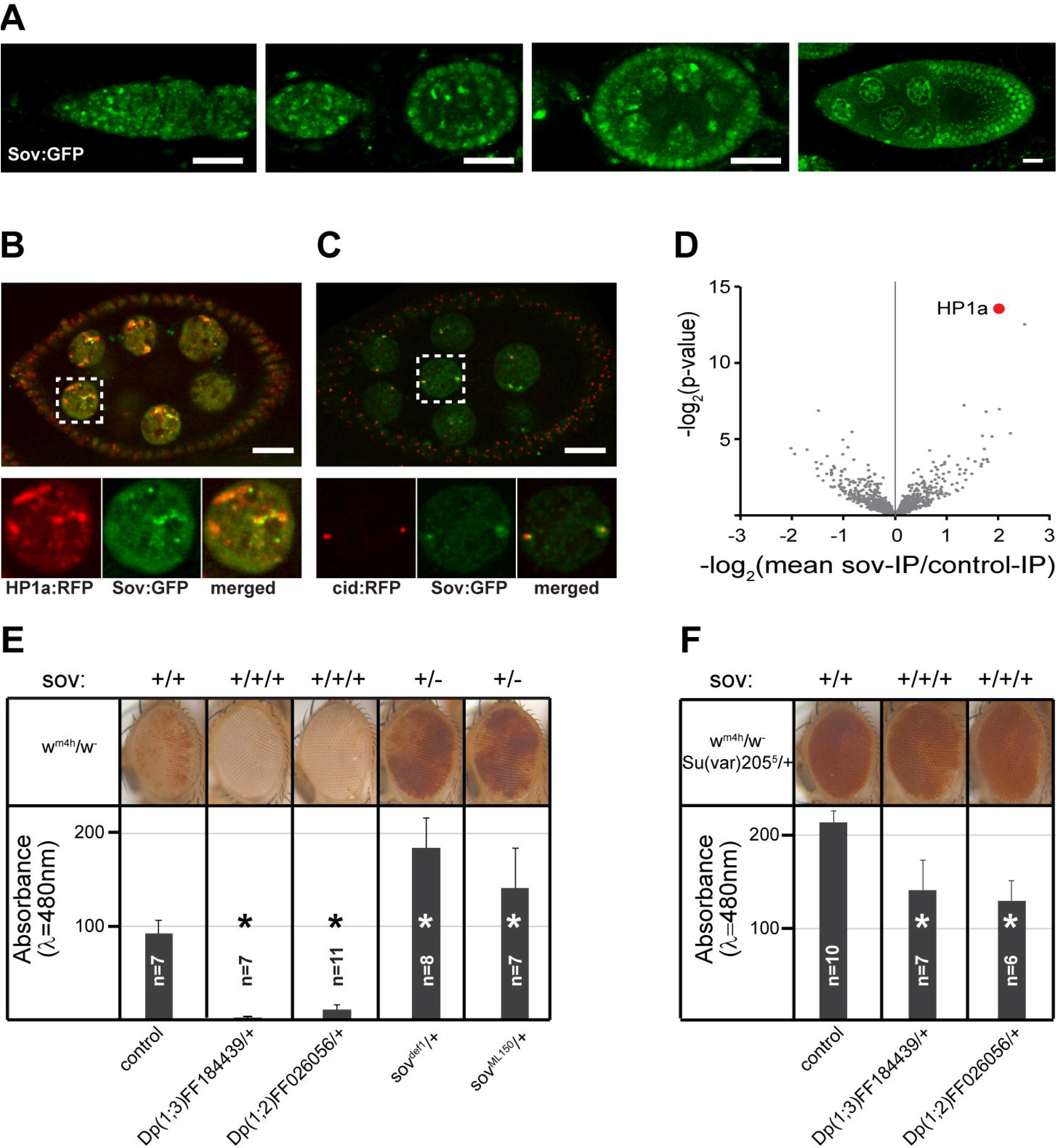
Sov interacts with HP1a and promotes heterochromatin formation. (A) Immunostainings showing localization of Sov:GFP at various stages of the oogenesis. Bars: 20μm. (B,C) Displayed are living egg chambers co-expressing Sov:GFP (green) with HP1a:RFP (red) in (B) and Cid:RFP (red) in (C), respectively. (D) Volcano plot showing fold enrichment values and significance levels for proteins co-immunoprecipitated with Sov:GFP from ovary lysates. HP1a (red dot) is the most significantly enriched protein. (E,F) Analysis of PEV modification effect of *sov.* Displayed are adult eyes and eye pigment levels of *w^m4h^/w^1118^* (E) and *w^m4h^/w^1118^; Su(var)205^5^/+* (F) females carrying various numbers of wild type copies of the *sov* gene. Mean±s.d. is shown, T-test, * pρ0.05

In summary, derepression of the reporter expression from the sensors and upregulation of the endogenous transposons by impaired *sov* function revealed an essential role for *sov* in *piwi-*mediated transposon repression, both in the germline and in the soma.

### Loss of *sov* induces Chk2-dependent checkpoint activation

Derepression of transposons results in accumulation of double strand breaks (DSBs) on the chromosomes, which are recognized by the histone variant, γH2Av. In the wild type, gH2Av accumulates in nuclei of germ cells in the 16 cell cysts undergoing meiotic recombination in region 2 of the germaria. As meiosis is accomplished, meiotic DSBs are repaired and γH2Av levels are reduced in the wild type germ cells (Fig.S3A). In contrast, an overall high level of γH2Av was detected in the nuclei of *sov* RNAi germ cells prior and after meiosis (Fig.S3B,C,D). Accumulation of γH2Av was initiated in the GSCs, indicating the formation of DSBs, presumably caused by transposon mobilization.

Transposon-induced DNA damage has been shown to activate the Chk2-dependent checkpoint. Activation of the Chk2 and ATR kinases induces microtubule organization defects in the oocytes, which lead to *gurken* RNA (grk), *oskar* RNA *(osk)* and Vasa delocalization and, thus, abnormal axis specification. In *sov* RNAi germ cells, no Vasa was detected at the posterior pole of the oocytes, indicating a disrupted AP polarity (Fig.S3E,F). Prior to Vasa, *osk* mRNA is localized to the posterior pole of the wild type oocytes where it is translated. However, neither *osk* mRNA (Fig.S3G,H) nor Osk protein (Fig.S3E,F) accumulation was detected at the posterior of *sov* depleted oocytes. In addition, no anterodorsal localization of Grk was detected in *sov* RNAi oocytes (Fig.S3E,F). Consistent with the lack of *Grk* localization, all eggs laid by *nosGal4>sovRNAi* females exhibited severe ventralized eggshell patterning defects with fused or absence of dorsal appendages (Fig.S3I,J). Analysis of the RNA-seq data revealed an increased expression of the apoptotic genes *hid* and *reaper* in the *sov* RNAi germ cells, indicating that the activation of the Chk2-dependent checkpoint is accompanied by induction of apoptosis (Fig.S3K).

Taken together, our data indicate that loss of *sov* function results in transposon derepression, which, in turn, leads to DNA damage and activation of the Chk2-dependent checkpoint.

### Sov promotes heterochromatin formation by stabilization of heterochromatic domains

To identify the mechanisms by which Sov regulates transposon repression, we first determined the subcellular localization GFP-tagged Sov expressed from a genomic fosmid construct. The tagged Sov variant completely rescued the sterile and the lethal phenotypes associated with *sov* mutations. In *sov^de11^; Sov:GFP* females, we detected ubiquitous Sov expression in the somatic and germ cells of the ovary. Sov localized in the nuclei and accumulated at nuclear foci (Fig.5A). At these foci, Sov partially co-localized with HP1a and Centrosome identifier (Cid), suggesting a direct role for Sov in chromatin regulation (Fig.5B,C).

To identify Sov interacting proteins, we affinity purified Sov from the *sov^del1^; Sov:GFP* ovaries on six independent occasions and performed mass spectrometry (LC-MS/MS) analysis of the samples. Consistent with the co-localization in the nuclei, Sov co-immunoprecipitated with HP1a, indicating that *sov* affects transposon repression as a chromatin regulator (Fig.5D). This role of Sov was confirmed in a pericentromeric position effect variegation (PEV)-assay. For the assay, the *w^m4h^* allele was used, in which the white gene responsible for the red eye color is translocated to the border between the pericentromeric heterochromatin and the euchromatin. In the individual facettes of the compound fly eye, stochastic spreading of the heterochromatin permits or inhibits expression of the white gene, leading to a variegating eye color (Elgin and Reuter, 2013).

In heterozygous *sov^ML150^* and *sov^del1^* flies, we detected increased eye pigment production, indicating a dominant suppressor effect of *sov* on PEV (Fig.5E). Dominant PEV-suppression of the *w^m4h^* allele suggests that Sov promotes heterochromatin formation at the pericentromeric regions of the genome. The positive effect of *sov* on heterochromatin formation is gene-dose-dependent, as demonstrated by the enhancement of PEV with increasing gene copies of *sov* (Fig.5E). Furthermore, PEV analysis revealed a dominant genetic interaction between HP1a (encoded by the *Su(var)205* gene) and *sov.* Increasing the copies of the wild type *sov* gene in *Su(var)205^5^* heterozygous flies caused decreased eye pigment production, indicating that increased amounts of Sov can compensate the heterochromatin formation defects caused by decreased HP1a levels (Fig.5F). Haplo-suppression and triplo-enhancement of PEV by *sov* and its genetic and physical interaction with HP1a indicates that Sov functions as a structural component of the heterochromatin.

Heterochromatin is epigenetically defined by repressive chromatin modifications, such as trimethylation of Histon3 at Lysin9 (H3K9me3), which is mediated by the histone methyl transferase *egg* and is recognized by HP1a. To test the effect of *sov* on the deposition of this repressive chromatin modification, we visualized H3K9me3 by immunostaining of *sov^del1^* homozygous germline clones. In the *sov* null mutant cells, no difference in H3K9me3 immunostaining was detectable when compared with the neighboring *sov* heterozygous cells, indicating that Sov functions downstream of *egg* in heterochromatin formation (Fig.S4A). Similar to the wild type, HP1a was recruited to the heterochromatic foci and formed elongated structures in *sov* homozygous germline clones and in *sov* RNAi germ cells (Fig.S4B-D). To dissect further the epistatic relationship between HP1a and sov, we analyzed Sov localization in HP1a silenced germ cells. Wild type Sov localization was found in HP1a RNAi cells, indicating that Sov localization does not depend on HP1a (Fig.S4E,F).

To test the involvement of Sov in HP1a mediated gene silencing, we applied a LacI/LacO transcriptional reporter assay. The NLS-GFP reporter construct containing LacO repeats was expressed in the ovary (Fig.6A,D). We expressed LacI tagged HP1a in the germline (HP1a:LacI) which was artificially recruited to the reporter via LacI-LacO interaction. Tethering HP1a:LacI to the reporter DNA suppressed GFP expression in the germline (Fig.6B,D). We silenced *sov* in the germline and observed no derepression of the reporter expression, suggesting that *sov* is not required for HP1a-mediated repression of the reporter locus when HP1a is artificially tethered to the DNA, but rather promotes the recruitment of HP1a to the chromatin. (Fig.6C,D).

**Fig. 6.**
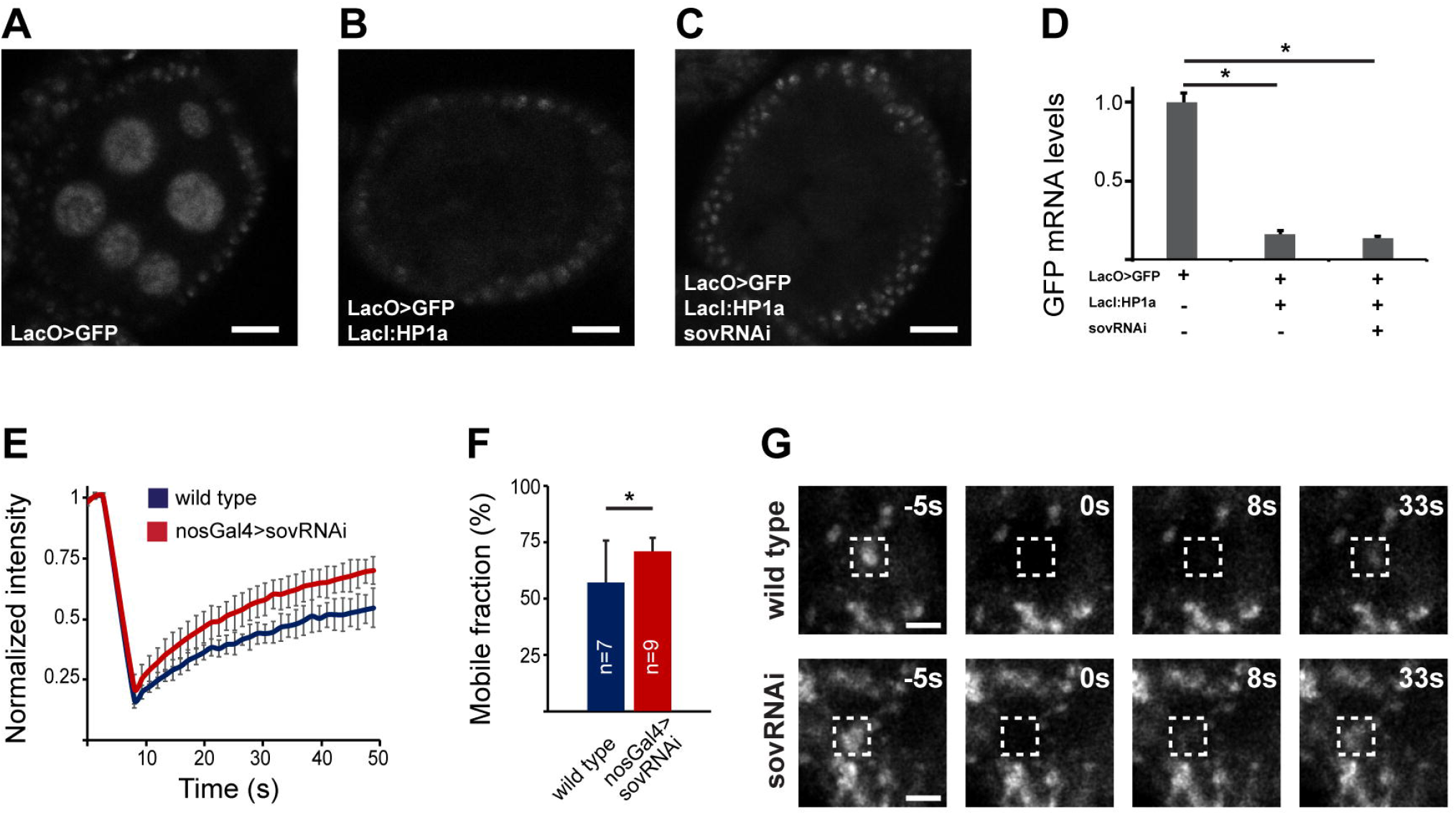
Sov stabilizes heterochromatin by the enhancement of HP1a association with the chromatin. (A-D) HP1a-tethering assay. (A) GFP fluorescence in egg chambers expressing the ubiquitous LacO-GFP reporter (A), the LacI:HP1a transgene (B) and the sovRNAi silencing construct (C) in the germ line. (D) Displayed are changes in GFP mRNA levels in ovaries expressing the indicated transgenes. Mean±s.d. is shown, T-test, * p<0.05. (E-G) Frap analysis of HP1a. Graphs showing fluorescence recovery curves (E) and mobile fractions (F) of HP1a:EGFP in wild type and sov-silenced germ cell nuclei. Mean±s.d. is shown, T-test, * p<.05 (G) Movie frames showing recovery of HP1a:EGFP fluorescence in wild-type and nosGal4>sovRNAi nuclei in a representative FRAP experiment. White boxes indicate photobleached regions. Bars: 2 μm.

Formation of heterochromatic domains have been shown to be driven by liquid phase separation via weak hydrophobic interaction between HP1a molecules and other heterochromatin components (Larson et al., 2017; Strom et al., 2017) In the mature heterochromatin, HP1a is mobile in the liquid compartment, whereas chromatin bound HP1a is immobile. To study the function of *sov* in the regulation of the dynamic properties of heterochromatin, we measured the immobile fraction of HP1a at the heterochromatin foci. We expressed HP1a: GFP in the germ cells and used fluorescent recovery after photobleaching (FRAP) to analyze HP1a dynamics (Movie1). Silencing of *sov* resulted in an increase of the mobile fraction in *sov* silenced germ cell nuclei (Fig.6E-G). Increase of HP1a mobility indicates that *sov* stabilizes the heterochromatin domain by the enhancement of HP1a association with the chromatin polymer.

### Sov promotes transcription in the heterochromatic genome regions

To determine the global effect of *sov* on steady state mRNA levels, we compared mRNA-seq data of *nosGal4>sovRNAi* and *nosGal4>wRNAi* control ovaries. Of the 6,811 euchromatic genes expressed in the ovary, transcription was activated by more than two-fold at 161 genes (2.4%), whereas 146 genes had a more than two-fold decrease in mRNA levels (2.1%). Of note is that of the 203 expressed genes mapping to the heterochromatic regions of the genome, 26 were downregulated more than two-fold (15.1%), whereas no heterochromatic gene was upregulated (Fig.7A). Over-representation of the heterochromatic genes in the downregulated gene set indicates that Sov preferentially promotes transcription in the heterochromatic genome regions.

**Fig. 7.**
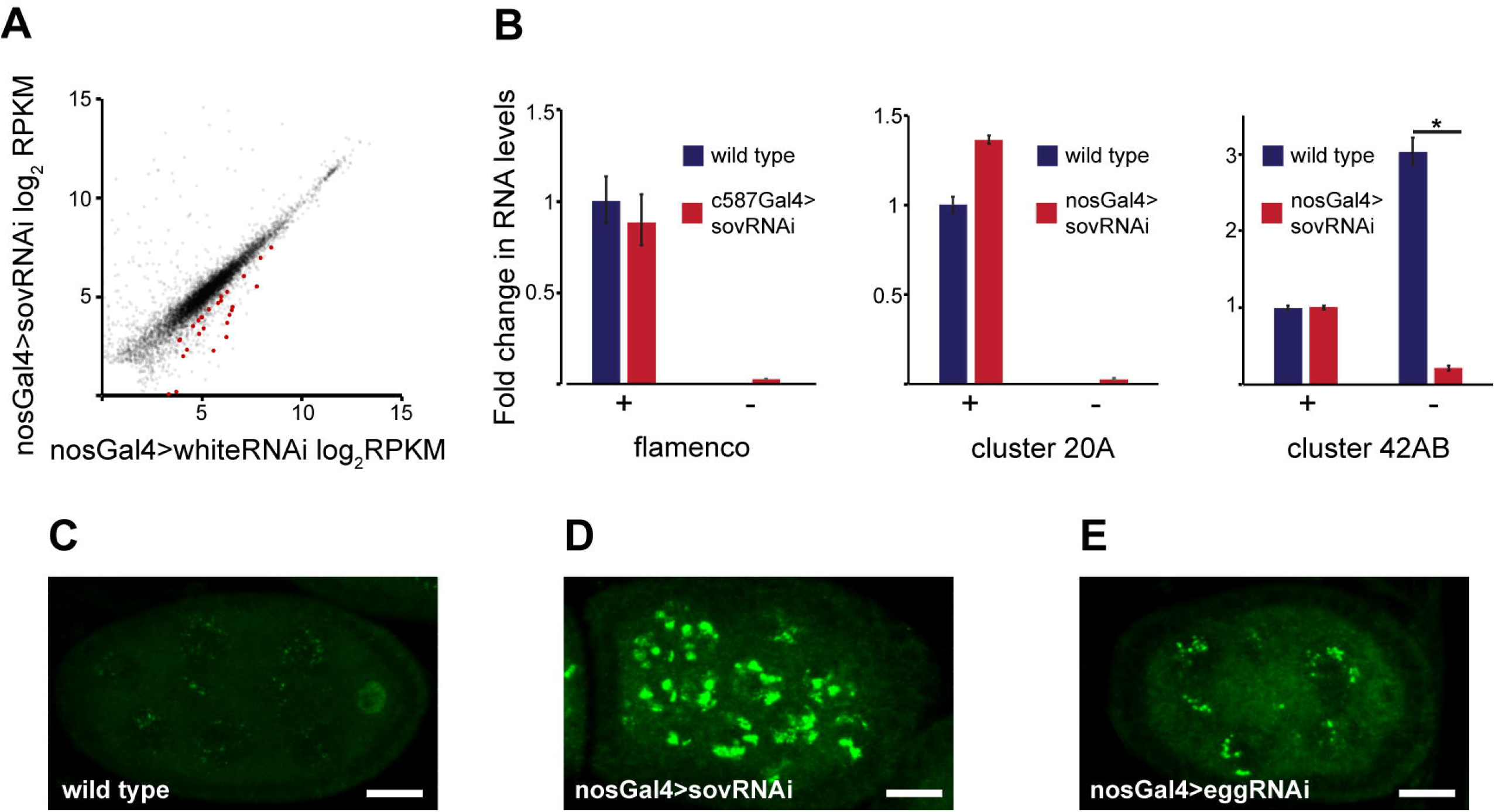
Sov promotes transcription in the heterochromatic genome regions. (A) Scatter plot showing expression levels of genes (gray) in nosGal4>sovRNAi ovaries. Heterochromatic genes with FC>2 are in shown in red. (B) Strand-specific RT-qPCR on wild type for RNAs derived from flamenco, cluster20A and cluster42AB. Mean±s.d. is shown, T-test, * pρ0.05. (C-E) Rhino (green) localization in wild type (C), sov-silenced (B), and *egg-*silenced (E) egg chambers.

Long piRNA precursors are transcribed from piRNA clusters which require a heterochromatic context for transcription. We hypothesized that *sov* is involved in transposon repression by regulating piRNA cluster transcription. In the soma, piRNA clusters are uni-strand clusters, i.e. only the plus strand of the piRNA clusters is transcribed. In the germ cells, piRNA precursors are generated from both uni-strand and dual-strand clusters. We carried out strand-specific RT-qPCR for piRNA precursors derived from the germline-specific dual-strand 42AB cluster, from the germline and soma-expressed uni-strand 20A cluster, and from the soma-specific *flam* cluster. In *sov* RNAi cells, no decrease in the transcript levels were detected from the transcribed strands of cl20A cluster and *flam*; however, transcription of the dual strand 42AB piRNA cluster was severely affected (Fig.7B). Although the plus-end specific transcript levels were not reduced, we detected decreased piRNA precursor production from the minus strand of the 42AB cluster. The effect of *sov* on transcription of the minus strand of the dual-strand piRNA cluster resembles that of *egg*, suggesting that *sov* affects chromatin regulation of the cluster (Rangan et al., 2011).

To investigate the chromatin structure at the dual-strand clusters, we analyzed the localization Rhino, an HP1 homologue associating exclusively with the dual-strand piRNA clusters. Silencing of *sov* induced formation of nuclear aggregates of Rhino suggesting that Sov is required for the formation of the proper chromatin structure at the dual-strand piRNA clusters (Fig.7C,D). Silencing of *egg* in the germline resulted in a similar abnormality in Rhino localization, supporting the hypothesis that Sov affects dual-strand cluster transcription by regulating heterochromatin structure at this cluster in cooperation with *egg* (Fig.7E).

Next, we analyzed whether *sov* silencing in the germ line affects expression of the genes involved in the piRNA pathway. Analysis of the RNA-seq data of germline-silenced *sov* ovaries revealed a significant downregulation of *Ago3,* a member of the Ago family required for piRNA biogenesis (Fig.S5A). Since mature piRNAs enable Piwi to enter into the nucleus, we analysed Piwi localisation in the *sov^del1^* mutant and in the *sov* RNAi cells. We detected a nuclear accumulation of Piwi, indicating that the nuclear translocation of Piwi is not affected by the reduction in *Ago3* levels (Fig.S6A-F). Nevertheless, reduction of *Ago3* expression in Sov-depleted germ cells uncovers the involvement of *sov* in an additional regulatory level of the piRNA-pathway.

## Discussion

### Sov is a novel heterochromatin component

Heterochromatin represents a functionally and structurally separate nuclear domain. The heterochromatin-associated DNA includes at least one-third of the Drosophila genome and is mainly composed of repetitive sequences, such as transposon fragments and satellite repeats (Hoskins et al., 2007) (Smith et al., 2007). Besides its DNA content, the heterochromatin is rather epigenetically defined by a pattern of various histone modifications and by the binding of additional chromatin-associated proteins (Riddle et al., 2011) (Saha et al., 2018) (Kharchenko et al., 2011). Although there is a diversity of combinatorial patterns of epigenetic marks in the heterochromatin, the majority of the heterochromatin regions lack the “activating” chromatin marks and are enriched in the “repressive” signatures, such as H3K9me3 and HP1a (Riddle et al., 2011). In addition to the DNA and the specific protein composition, heterochromatin has a higher level of organisation. HP1a is capable of demixing from solution and forming liquid droplets that organize the assembly of the heterochromatin into a membrane-less nuclear domain via liquid-liquid phase separation (Larson et al., 2017; Strom et al., 2017). Many proteins driving the formation of phase-separated organelles contain extended intrinsically disordered regions and are enriched in RGG/RG domains. Sov shows a remarkable structural similarity to these proteins since it has an unusually long disordered N-terminus that contains an RGG/RG repeat domain. It is tempting to speculate that Sov functions, in concert with its binding partner HP1a, in the establishment of the special physical properties of the heterochromatin. Indeed, the integrity of the mature heterochromatin domain was shown to require formation of the immobile HP1a compartment through the interaction of HP1a with non-histone binding partners (Strom et al., 2017). We propose that Sov is one of these non-histone partners of HP1a. We show that Sov affects the dynamics of HP1a between the liquid and static compartments in the heterochromatin domain, suggesting that the HP1a-mediated phase separation driven heterochromatin formation depends, at least in part, on Sov. The RGG/RG domain found in Sov is a widespread RNA binding domain displaying degenerate RNA binding specificity (Ozdilek et al., 2017). The HP1a complex containing Sov is enriched in RNAs, raising the possibility that the putative RNA binding of Sov is involved in proper heterochromatin function (Alekseyenko et al.,2014). Targeting of heterochromatin formation at particular genomic regions involves diverse mechanisms, such as function of satellite DNA-specific binding proteins or the RNAi machinery (Elgin and Reuter, 2013). In the LacI/LacO-based tethering assay, we show that HP1a-mediated repression is independent of Sov, while Sov enhances the recruitment and binding of HP1 to the chromatin. This function of Sov could be performed by the highly structured C-terminus, which contains a large number of tandem Zn fingers. Through these domains, Sov may stabilize the binding of HP1a complexes to specific DNA sequences and target Sov/HP1a-mediated processes at specific regions of the genome.

Heterochromatin is usually associated with transcriptional repression (Smith et al., 2007). Many protein-coding genes residing in the repressive domains, however, require the heterochromatic environment for transcription (Clegg et al., 1998; Lu et al., 2000; Schulze et al., 2005; Wakimoto and Hearn, 1990). Disruption of heterochromatin by impaired HP1a function results in reduced expression of these genes (Lundberg et al., 2013). Consistent with this observation, Sov preferentially promotes the transcription of heterochromatin-resident genes by positively regulating HP1 function.

### Sov is involved in the piRNA pathway

We demonstrate that Sov is involved in the regulation of the piRNA pathway at multiple levels. In the germline, it is required for proper transcription of the piRNA clusters and promotes the expression of the piRNA-pathway component, Ago3. In addition, it may mediate transcriptional gene silencing, both in the germline and in the somatic cells.

Efficient transcription of the piRNA clusters requires a heterochromatic environment. Despite the obvious differences in their regulation, both uni-strand and dual-strand clusters were shown to be enriched in the repressive H3k9m3 histone mark. Impairment of heterochromatin formation leads to reduction of piRNA cluster transcription. (Rangan et al., 2011) (Teo et al., 2018). The transcription of the uni-strand clusters critically depends on *egg,* however; exclusively the minus strand of the dual strand 42AB cluster was sensitive to *egg* depletion (Rangan et al., 2011). Similar to *egg,* depletion of *sov* results in reduction of the transcription of the minus strand of the 42AB cluster, while the transcription of the plus strand remains unaffected. The effect of *egg* and *sov* depletion on cluster transcription, however, is not identical. In contrast to *egg,* the function of *sov* is restricted to the dual strand cluster. It is possible that *sov* specifically regulates the formation of the Rhino-dependent specialized heterochromatin that enables dual-strand transcription. Further studies are needed to explore how this separation in *sov* function is regulated.

Another level of transposon regulation, where Sov is involved, is the indirect control of piRNA biogenesis. Like *egg, sov* is required for *Ago3* transcription, raising the possibility that the transposon derepression defects found in *sov* depleted germ cells may result not only from reduced cluster transcription, but also from defective piRNA biogenesis (Kang et al., 2018). However, the effect of *sov* on transposon repression and niche regulation differs from that of *Ago3.* Unlike *egg* or *sov* mutants, no GSC loss was detected in the *Ago3* mutants, showing that GSC self-renewal was not affected (Rojas-Ríos et al., 2017). Consistent with the lack of GSC self-renewal defect in *Ago3* mutants, piRNAs are produced in the absence of Ago3 through an Aub-dependent homotypic amplification mechanism, and these piRNAs could be used in the regulation of GSCs (Wang et al., 2015a). piRNAs enable Piwi to move from the cytoplasm to the nucleus, where it suppresses transposon transcription (Wang and Elgin,2011; Klenov et al., 2011; Sienski et al., 2012; Rozhkov et al.,2013; Le Thomas et al., 2013) (Yashiro et al., 2018). In *sov* mutant and sov RNAi cells, we detect nuclear Piwi accumulation, suggesting that piRNA biogenesis was not completely abolished. Loss of Ago3 results in the derepression of a subset of transposons (Wang et al., 2015a). Sov depletion, however, causes a stronger defect in transposon regulation, i.e. the derepression of all transposon classes, indicating that *sov* has a more general effect on transposon silencing than the sole enhancement of *Ago3* transcription.

In the *sov* mutant ECs, we find a derepression of the *gypsy* transposon. piRNAs required for *gypsy* silencing are generated from the *flam* cluster. Similar to HP1a, Sov is dispensable for *flam* transcription (Penke et al., 2016). Since there is no Ago3-dependent piRNA amplification in the soma, Sov must have an additional effect on transposon silencing. An attractive explanation for *gypsy* derepression in the Sov-depleted ECs could be that *sov* is involved in the transcriptional transposon silencing mechanism. This step of the silencing pathway requires heterochromatinization of the transposon locus. The initiation of this process is the targeting of the nuclear piwi-RISC to the nascent transposon transcripts based on the complementarity between the transposon mRNA and the Piwi-bound piRNA. Tethering of the Piwi-RISC to the transposon induces the recruitment of the effector complex, composed of Asterix, Panx, Mael, and Egg, to the transposon loci (Sato and Siomi, 2018). Egg initiates the deposition of the repressive H3K9me3 mark and inhibits transcription by recruiting HP1a. In a parallel pathway, Piwi also recruits Histon1 to the transposons, which organizes the chromatin into a higher order repressive state (Iwasaki et al., 2016). Sov physically interacts with HP1a and stabilizes the heterochromatin domain through the enhancement of HP1a association with the chromatin. We propose that Sov supports the recruitment of HP1a to the transposon locus by binding to HP1a, which in turn stabilizes the association of H3K9me3-bound HP1a with the chromatin. A similar Sov-mediated mechanism can work also in the germline, however, this requires further investigation.

### Sov functions in the stem cell niche

We show that Sov is involved in germ cell development by regulating GSC survival, GSC differentiation and EC survival. The complex loss-of-function phenotype of *sov* closely resembles that of *egg* mutations, supporting our hypothesis that *sov* contributes to the formation and maintenance of the Hp1a-Egg mediated repressive chromatin environment.

Germline-specific RNAi revealed that Sov is required in the GSCs to control their self-renewal in a cell-autonomous manner. Loss of function of the components of the heterochromatin machinery and of the piRNA pathway induce apoptotic GSC loss accompanied with transposon derepression, DNA damage and checkpoint activation. (Ma et al., 2014; Ma et al., 2017). In *egg* mutants, upregulation of apoptotic genes *hid, reaper,* and activation of Caspase3 cleavage was found (Clough et al., 2007; Kang et al., 2018). Similarly, we observed an accumulation of DNA damage, checkpoint activation and apoptosis in *sov*-depleted cells. We propose that *sov* ensures GSC maintenance through suppression of transposon activity induced genome damage and by so doing, supresses the apoptosis of the germ cells.

Cell-type specific RNAi experiments showed that Sov function is required in EC cells for controlling both the maintenance and the differentiation of germ cells in a non-cell-autonomous manner. When anterior-most ECs are lost, GSCs are gradually eliminated from the niche, while loss of posterior ECs leads to differentiation defects and to the formation of GSC tumors (Wang and Page-McCaw, 2018). Indeed, EC-specific *sov* RNAi resulted in loss of both the anterior and the posterior ECs and, as a consequence, a combination of the GSC loss and germ cell tumour phenotypes was observed.

The *sov* dependent EC loss may be induced by the improper function of the signalling network operating in the somatic niche. We show that *sov* has an effect on the *Wnt* and *dpp* signalling pathways. *Wnt* signalling was shown to promote survival of ECs and inhibit the expansion of *dpp* activity in the niche (Wang et al., 2011; Wang et al., 2015b). Interestingly, transposon derepression results in decreased Wnt4 expression (Upadhyay et al., 2016). We show that *sov* is required for Wnt4 expression in the ECs, however, the differentiation defect induced by *sov* depletion is not rescued by simultaneous Wnt overexpression indicating additional diverse roles of *sov* in ensuring EC survival.

In ECs, increasing transposon activity by knocking down egg, *hp1a, piwi* or *flam* results in cell death (Upadhyay et al., 2016; Wang et al., 2011) (Ma et al., 2014). Since in *sov* depleted ECs, we detected robust transposon derepression accompanied by caspase activation, we propose that Sov supresses EC death by down-regulating transposons.

## Materials and methods

### Drosophila stocks

All animals were raised at 25°C, unless otherwise indicated. The *sov^de11^* allele was generated by FRT-mediated recombination between PBac{WH}f04480 and P{XP}d07849 transposon insertions (Ryder et al., 2004). The *sov^2^* and *sov^ML150^* alleles were obtained from the Bloomington Drosophila stock center. To silence sov, we generated the *UAS-sovRNAi[3-2]* transgenic construct expressing shRNA targeting the last exon of *sov*. In this study, *UAS-sovRNAi[3-2]* construct was used for *sov* silencing unless otherwise indicated. The Dp(1;3)FF184439 and Dp(1;2)FF026056 duplications were generated by inserting the FF184439 and the FF026056 fosmid constructs into the attP2 and attP40 docking sites, respectively (Ejsmont et al., 2011). The FlyFos018439-CG14438-SGFP-V5-preTEV-BLRP-3XFLAG fosmid was obtained from the Drosophila TransgeneOme Project and was inserted into the attP40 site. For the sake of simplicity, we refer this transgenic line Sov:GFP thereafter.

The strains used in this study include: *sov^2^* (Bl#4611), *sov^ML150^* (Bl#4591), *bam-GFP* (Chen and McKearin, 2003), *nos-Gal4* (Van Doren et al., 1998), *c587Gal4* (Manseau et al., 1997; Song et al., 2004), *hs-bam* (Ohlstein and McKearin, 1997), *gypsy-LacZ* (Dönertas et al., 2013), *Burdock-LacZ* (Dönertas et al., 2013), *Het-A-LacZ* (Dönertas et al., 2013), *Vasa:AID:GFP*(Bl# 76126) (Bence et al., 2017), *HP1a:RFP*(BL#30562) (Wen et al., 2008), *HP1a:GFP* (BL#30561) (Wen et al., 2008), *FRT19A,His2Av:GFP,hsFLP/FRT19A; His2Av-mRFP* (B#30563), FRT19A Bl#1709), *Su(var)205^5^* (Bl#6234), *w^m4h^, UAS-HP1aRNAi-TRiP.GL00531* (Bl#36792), *UAS-eggRNAi-TRiP.HMS00443* (Bl#32445); *UAS-wRNAi-TRiP .HMS0001* (Bl#33623); *UAS-sovRNAi-TRiP.HMC04875* (Bl#57558), *UAS-sovRNAi-KK103679* (VDRC#v100109), *8XLacO-nos>GFP* (Sienski et al., 2015), *UASP-LacI:HP1a* (Sienski et al., 2015) *UAS-Wnt4.ORF.3XHA* (FlyORF#F001112), *UAS-dWnt4(r13)* (Sato et al., 2006), *oskMS2/MS2GFP*(Forrest and Gavis, 2003a; Zimyanin et al., 2008).

### Sequencing of the *sov^2^* mutant allele

To sequence the *sov^2^* mutant allele, genomic DNA was isolated from *sov^2^/sov^del1^* mutant females and the *sov* coding sequence, including the introns, was PCR-amplified and sequenced between the start and the stop codons. Comparison of the *sov* sequence with the reference R6.15 Drosophila genomic sequence revealed 20 missense mutations, a 27 bp-long in-frame deletion, a 54 bp-long in frame insertion and a frame-shift mutation in the *sov^2^* coding region. To unambiguously identify the mutation responsible for the *sov* mutant phenotype, we sequenced the *sov* coding sequence of the *wisp^12^’^3147^* allele and used it as a reference. The *wisp^12-3147^* and the *sov^2^* alleles were isolated in the same genetic screen and have the same paternal chromosome. Analysis of the sequences revealed that all mutations are shared between *sov^2^* and *wisp^12^’^3147^,* except for the frame-shift mutation.

### Clone induction and cell type-specific silencing

The marked control and *sov* mutant EC and germ line clones were generated using the FLP/FRT-mediated recombination as described in (Song et al., 2002; Xie and Spradling, 1998). To generate *sov* mutant clones, *FRT19A,His2Av:GFP,hsFLP/FRT19A,sov^del1^; His2Av-mRFP/+* females were heat shocked either 24 hours after egg laying or at the adult stage. Young females were heat shock on three consecutive days for one hour at 37°C, and the mutant phenotypes were examined six days after clone induction. To silence *sov* in the ECs specifically at the adult stage, *c587Gal4UAS-sovRNAi[3-2];TubGal80^ts^* females were cultured at 18°C until adulthood, and they were then shifted to 29°C. For the transposon sensor assay, *sov* was silenced using the germ line specific MTDGal4 driver. To express Bam in *sov* mutants, *sov^2^/sov^ML150^; hs-bam* flies were heat-shocked at 37°C two times for one hour, separated by a two hour recovery period at 25°C. Ovaries were analyzed 24 hours after the first heat-shock.

### Position effect variegation assay

Female flies were aged at 25°C for 10 days prior to imaging. To measure eye pigmentation, females were frozen in liquid nitrogen. Heads were separated from bodies by brief vortexing. Samples of 10 females were homogenized in 0.5 ml of 0.01M HCl in ethanol; the homogenate was placed at 4°C overnight, warmed at 50°C for 5 min, clarified by centrifugation, and the OD at 480 nm of the supernatant was measured.

### Immunohistochemistry and FRAP

β-Gal staining of transposon sensor lines was performed as described in (Dönertas et al., 2013) Immunostainings were performed as described earlier (Jankovics et al., 2014). List of primary antibodies used is summarized in TableS1. DAPI was used to label the nuclei, actin was visualized by phalloidin staining. Specimens were examined with Leica TCS SP5 confocal microscope. For live imaging of Sov:GFP, HP1a:GFP, HP1a:RFP, and Cid:RFP, samples were prepared as described in (Forrest and Gavis, 2003b) and examined with VisiScope spinning disc confocal microscope. Fluorescence recovery experiments were performed on stage 10 egg chambers expressing HP1a:GFP. FRAP experiments were performed with a Leica SP5 confocal microscope. The 405 nm laser line was used to photobleach a 4μm^2^ region of the heterochromatin domain. Recovery was recorded for one minute at 1 frame every 1.5s. Fluorescence recovery curves were analyzed using the easyFrap software as described in (Bancaud et al., 2010; Koulouras et al., 2018). Statistical tests were performed with GraphPad Prism.

### Co-immunoprecipitation and mass spectrometry

For the protein interactome analysis, two-day old *Sov:GFP* and *w^1118^* females were used. Ovaries were dissected in PBS. 150-200 μl of ovaries were harvested, frozen in liquid nitrogen and ground with TissueLyser at 30Hz. Total proteins from *Drosophila* tissue were extracted using the manufacturer’s Lysis buffer supplemented 1 mM DTT, 1 mM PMSF, 1xsigma protease inhibitor cocktail, 3 mM pNPP, 1 uM MG132. Total protein extracts (8mg/IP) were immunopurified using anti-GFP (three replicates) or anti-FLAG (three replicates) antibody coupled 50 nm magnetic beads (MACS Technology, Miltenyi) digested in column with trypsin, and analyzed in a single run on the mass spectrometer [Hubner et al. 2010]

The resulting peptide mixture was desalted prior to LC-MS/MS analysis (Omix C18 100 ul tips, Varian) and the purified peptide mixture was analyzed by LC-MS/MS using a nanoflow RP-HPLC (Lc program: linear gradient of 3-40 % B in 100 min, solvent A: 0.1% formic acid in water, solvent B: 0.1% formic acid in acetonitrile) on-line coupled to a linear ion trap-Orbitrap (Orbitrap-Elite, Thermo Fisher Scientific) mass spectrometer operating in positive ion mode. Data acquisition was carried out in data-dependent fashion, the 10 most abundant, multiply charged ions were selected from each MS survey for MS/MS analysis (MS spectra were acquired in the Orbitrap, and CID spectra in the linear ion trap).

Raw data were converted into peak lists using the in-house PAVA software [Guan, S.; Price, J. C.; Prusiner, S. B.; Ghaemmaghami, S.; Burlingame, A. L. Mol Cell Proteomics 2011, 10, M111.010728 doi: 10.1074/mcp.M111.010728.] and searched against the Swissprot database (downloaded 2015/04/16, 548208 proteins) using the Protein Prospector search engine (v5.15.1) with the following parameters: enzyme: trypsin with maximum 1 missed cleavage; mass accuracies: 5 ppm for precursor ions and 0.6 Da for fragment ions (both monoisotopic); fixed modification: carbamidomethylation of Cys residues; variable modifications: acetylation of protein N-termini; Met oxidation; cyclization of N-terminal Gln residues allowing maximum 2 variable modifications per peptide. Acceptance criteria: minimum scores: 22 and 15; maximum E values: 0.01 and 0.05 for protein and peptide identifications, respectively. Another database search was also performed using the same search and acceptance parameters except that Uniprot. random. concat database (downloaded 2015/4/16) was searched with Drosophila melanogaster species restriction (52524 proteins) including additional proteins identified from the previous Swissprot search (protein score>50). False discovery rate was estimated using peptide identifications representing randomized proteins (2* #of random IDs/total peptide IDs) = 2 times number of random IDs divided by peptide IDs.

Spectral counting was used to estimate relative abundance of individual proteins in the SovGFP and control samples: peptide counts of the individual proteins were normalized to the total number of peptide identifications in each sample, then these normalized peptide counts were compared in the two samples. Enrichment between SovGFP and control experiments were calculated by edgeR (Li and Andrade, 2017). Counts representing Sov were omitted from the analysis.

### RT-quantitative PCR

In general, the total RNA was prepared from whole ovaries using ReliaPrep RNA Tissue Miniprep System (Promega, Z6111). cDNAs were synthetized using oligodT primers (First Strand cDNA Synthesis kit, ThermoScientific, K1612). Strand specific RT-qPCR was performed as described (Klattenhoff et al., 2009). For strand specific RT-qPCR of the 42AB and 20A piRNA clusters, ovaries of two-day old *nosGal4>sovRNAi* females were used. For strand specific RT-qPCR of the *flam* cluster, *c587Gal4>sovRNAi; Tub>Gal80^ts^* females were cultured on 18°C and shifted to 29°C at the adult stage. RT-qPCR was performed 10 days after the temperature shift. qPCR reactions were performed using Maxima SYBR Green/ROX (ThermoScientific, K0221). For each reaction, tree technical replicates were performed on three biological replicates. Rp49 transcript was used as internal control. List of primer sequences is shown in Supplementary table 2. Data ware analyzed using the Rotor-Gene Q Sereies software. Primer sequences are provided in Supplementary Table 2.

### RNA-sequencing and data analysis

For RNA-seq, ovaries of *nosGal4>sovRNAi3-2* and *nosGal4>wRNAi* females were dissected. Total RNA was prepared using RNeasy Mini Kit (Qiagen, #74104). RNA samples were quantified and their quality determined by capillary gel electrophoresis with a 2100 Bioanalyzer (Agilent) instrument using Agilent RNA 6000 nano kit. Poly(A) RNAs were selected with NEBNext Poly(A) mRNA Magnetic Isolation Module then strand-specific RNA-seq libraries were prepared using NEBNext Ultra RNA Library Prep Kit for Illumina with NEBNext Multiplex Oligos for Illumina following the recommendations of the manufacturer (New England Biolabs). Sequencing libraries were validated and quantified with Agilent DNA 1000 kit in a 2100 Bioanalyzer instrument. After pooling, paired-end sequencing was done with an Illumina MiSeq instrument using MiSeq Reagent kit V3-150. Fastq files were generated with MiSeq Control Software then quality trimmed and filtered with Trimmomatic v0.33. For transcriptome analysis sequence flies were aligned to the Drosophila melanogaster reference genome r6.13 with Tophat v2.0.9. After alignment files were deduplicated with SAMtools software and differential expression analysis was performed with CuffDiff v2.1.1 using the corresponding transcript annotation file. For transposon expression analysis trimmed fastq files were aligned to the USCS dm6 reference genome with Bowtie v2.1 then differential transposon expression was determined by CuffDiff v2.1.1 using a corresponding transposon annotation file (http://labshare.cshl.edu/shares/mhammelllab/www-data/TEToolkit/TE_GTF/) (Jin et al., 2015).

## Acknowledgements

We thank W. Theurkauf, T. Tetsuya, D. McKearin, L. Gilboa and J. Brennecke for fly stocks and reagents. We also thank B. Irvine for critical reading of the manuscript. This work was supported by grants from the National Research, Development and Innovation Office (NKFIHK112294, NKFIK117010, OTKA-K108538, GINOP-2.3.2-15-2016-00032, GINOP-2.3.2.-15-2016-00001). M.B. is supported by the János Bolyai Fellowship.

